# Full-genome evolutionary analysis of the novel corona virus (2019-nCoV) rejects the hypothesis of emergence as a result of a recent recombination event

**DOI:** 10.1101/2020.01.26.920249

**Authors:** D. Paraskevis, E.G. Kostaki, G. Magiorkinis, G. Panayiotakopoulos, S. Tsiodras

## Abstract

**Background:** A novel coronavirus (2019-nCoV) associated with human to human transmission and severe human infection has been recently reported from the city of Wuhan in China. Our objectives were to characterize the genetic relationships of the 2019-nCoV and to search for putative recombination within the subgenus of sarbecovirus.

**Methods:** Putative recombination was investigated by RDP4 and Simplot v3.5.1 and discordant phylogenetic clustering in individual genomic fragments was confirmed by phylogenetic analysis using maximum likelihood and Bayesian methods.

**Results:** Our analysis suggests that the 2019-nCoV although closely related to BatCoV RaTG13 sequence throughout the genome (sequence similarity 96.3%), shows discordant clustering with the Bat-SARS-like coronavirus sequences. Specifically, in the 5’-part spanning the first 11,498 nucleotides and the last 3’-part spanning 24,341-30,696 positions, 2019-nCoV and RaTG13 formed a single cluster with Bat-SARS-like coronavirus sequences, whereas in the middle region spanning the 3’-end of ORF1a, the ORF1b and almost half of the spike regions, 2019-nCoV and RaTG13 grouped in a separate distant lineage within the sarbecovirus branch.

**Conclusions:** The levels of genetic similarity between the 2019-nCoV and RaTG13 suggest that the latter does not provide the exact variant that caused the outbreak in humans, but the hypothesis that 2019-nCoV has originated from bats is very likely. We show evidence that the novel coronavirus (2019-nCov) is not-mosaic consisting in almost half of its genome of a distinct lineage within the betacoronavirus. These genomic features and their potential association with virus characteristics and virulence in humans need further attention.

## Main text

The family Coronaviridae includes a large number of viruses that in nature are found in fish, birds and mammals[1, 2]. Human coronaviruses, first characterized in the 1960s are associated with a large percentage of respiratory infections both in children and adults[1, 3].

Scientific interest in Coronaviruses exponentially increased after the emergence of SARS-Coronavirus (SARS-CoV) in Southern China[4–6]. Its rapid spread led to the global appearance of more than 8,000 human cases and 774 deaths[1]. The virus was initially detected in Himalyan palm civets[7] that may have served as an amplification host; the civet virus contained a 29-nucleotide sequence not found in most human isolates that were related to the global epidemic[7]. It has been speculated that the function of the affected open reading frame (ORF 10) might have played a role in the trans-species jump[1]. Later on a similar virus was found in horseshoe bats[8, 9]. A 29-bp insertion in ORF 8 of bat-SARS-CoV genome, not found in most human SARS-CoV genomes, was suggestive of a common ancestor with civet SARS-CoV [8]. After the SARS epidemic bats have been considered as a potential reservoir species that could be implicated in future coronavirus-related human pandemics [10]. During 2012 Middle East Respiratory coronavirus (MERS-CoV) emerged in Saudi Arabia [11, 12] and has since claimed the lives of 919 out of 2521 (35%) people affected [13]. A main role in the transmission of the virus to humans has been attributed to dromedary camels [14]and its origin has been again traced to bats [15].

Ever since both SARS and MERS-CoV (due to their high case fatality rates) are prioritized together with “highly pathogenic coronaviral diseases other than MERS and SARS” under the Research and Development Blueprint published by the WHO [16].

A novel coronavirus (2019-nCoV) associated with human to human transmission and severe human infection has been recently reported from the city of Wuhan in the Hubei province in China [17, 18]. A total of 1320 confirmed and 1965 suspect cases were reported up to 25 January 2020; of the confirmed cases 237 were severely ill and 41 had died [17]. Most of the original cases had close contact with a local fresh seafood and animal market [19, 20].

Full-genome sequence analysis of 2019-nCoV revealed that belongs to betacoronavirus, but it is divergent from SARS-CoV and MERS-CoV that caused epidemics in the past [19]. The 2019-nCoV together with the Bat_SARS-like coronavirus forms a distinct lineage within the subgenus of the sarbecovirus [19].

Our objectives were to characterize the genetic relationships of the 2019-nCoV and to search for putative recombination within the subgenus of sarbecovirus.

Viral sequences were downloaded from NCBI nucleotide sequence database (http://www.ncbi.nlm.nih.gov). The BatCoV RaTG13 sequence was downloaded from the GISAID BetaCov 2019-2020 repository (http://www.GISAID.org). The sequence was reported in Zhu et al [21]. Full-genomic sequence alignment was performed using MAFFT v7.4.2. [22] and manually edited using MEGA v1.0 [23] according to the encoded reading frame. Putative recombination was investigated by RDP4 [24] and Simplot v3.5.1 [25] and discordant phylogenetic clustering in individual genomic fragments was confirmed by phylogenetic analysis using maximum likelihood (ML) and Bayesian methods. ML trees were reconstructed using Neighbor-Joining (NJ) with ML distances or after heuristic ML search (TBR) with GTR+G as nucleotide substitution model as implemented in PAUP* 4.0 beta [26]. The GTR+G was used in Bayesian analysis as implemented in MrBayes v3.2.7. [27] Phylogenetic trees were viewed using FigTree v1.4. (http://tree.bio.ed.ac.uk/software/figtree/)

A similarity plot was performed using a sliding window of 450 nts moving in steps of 50 nts, between the query sequence (2019-nCoV) and different sequences grouped according to their clustering pattern. The similarity plot (figure 1A, B) suggested that the RaTG13 was the most closely related sequence to the 2019-nCoV throughout the genome. The genetic similarity between the 2019-nCoV and RaTG13 was 96.3% (p-uncorrected distance: 0.0369). On the other hand, a discordant relationship was detected between the query and the sequences of the Bat-SARS-like coronavirus (MG772934 and MG772933) (figure 1C). Specifically in in the 5’-part of the genome spanning the first 10,901 nts of the alignment that correspond to the 11,498 nucleotides of the prototype strain (NC_045512) and the last 3’-part spanning 22,831-27,933 positions (24,341-30,696 nucleotides in the NC_045512), 2019-nCoV and RaTG13 formed a single cluster with Bat_SARS-like coronavirus sequences (figure 1C). In the middle region spanning the 3’-end of ORF1a, the ORF1b and almost half of the spike regions (10,901-22,830 nts in the alignment or 11,499-24,340 of the NC_045512), 2019-nCoV and RaTG13 grouped in a separate distant lineage within the sarbecovirus branch (figure 1B, 1C). In this region the 2019-nCoV and RaTG13 is distantly related to the Bat_SARS-like coronavirus sequences. Phylogenetic analyses using different methods confirmed these findings. A BLAST search of 2019-nCoV middle fragment revealed no considerable similarity with any of the previously characterized corona viruses (figure 2).

**Figure 1.**
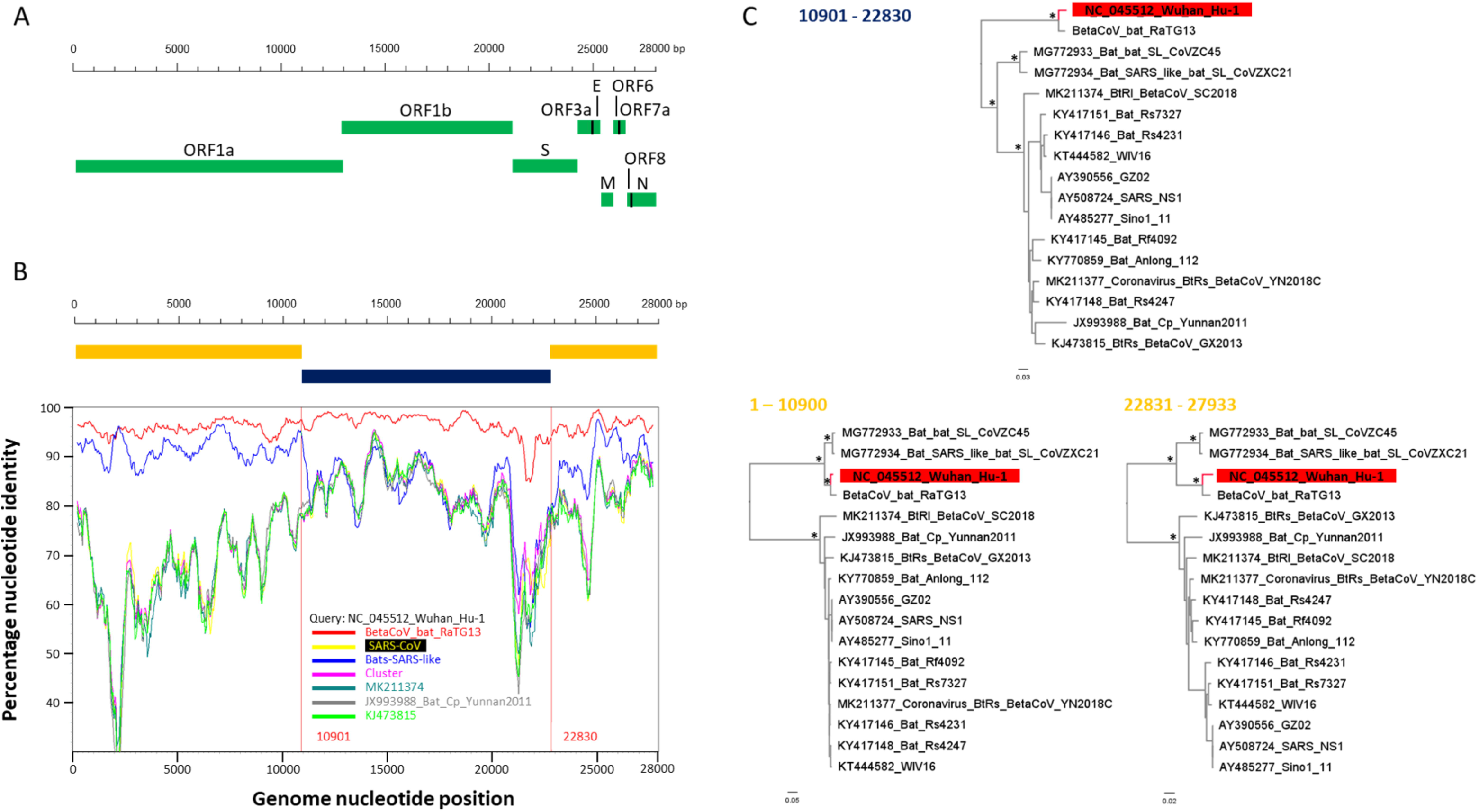
**A**. Genomic organization of the novel coronavirus (2019-nCoV) according to the positions in the edited alignment. **B**. Simplot of 2019-nCoV (NC_045512_Wuhan_Hu-1) against sequences within the subgenus sarbecovirus. Different colours correspond to the nucleotide similarity between the 2019-nCoV and different groups. The regions with discordant phylogenetic clustering of the 2019-nCoV with Bats-SARS-like sequences are shown in different colours. **C**. Maximum likelihood (ML) phylogenetic trees inferred in different genomic regions as indicated by the Simplot analysis. The genomic regions are shown in numbers at the top or at the left of the trees. The 2019-nCoV sequence is shown in red and stars indicate important nodes received 100% bootstrap and 1 posterior probability support.

**Figure 2.**
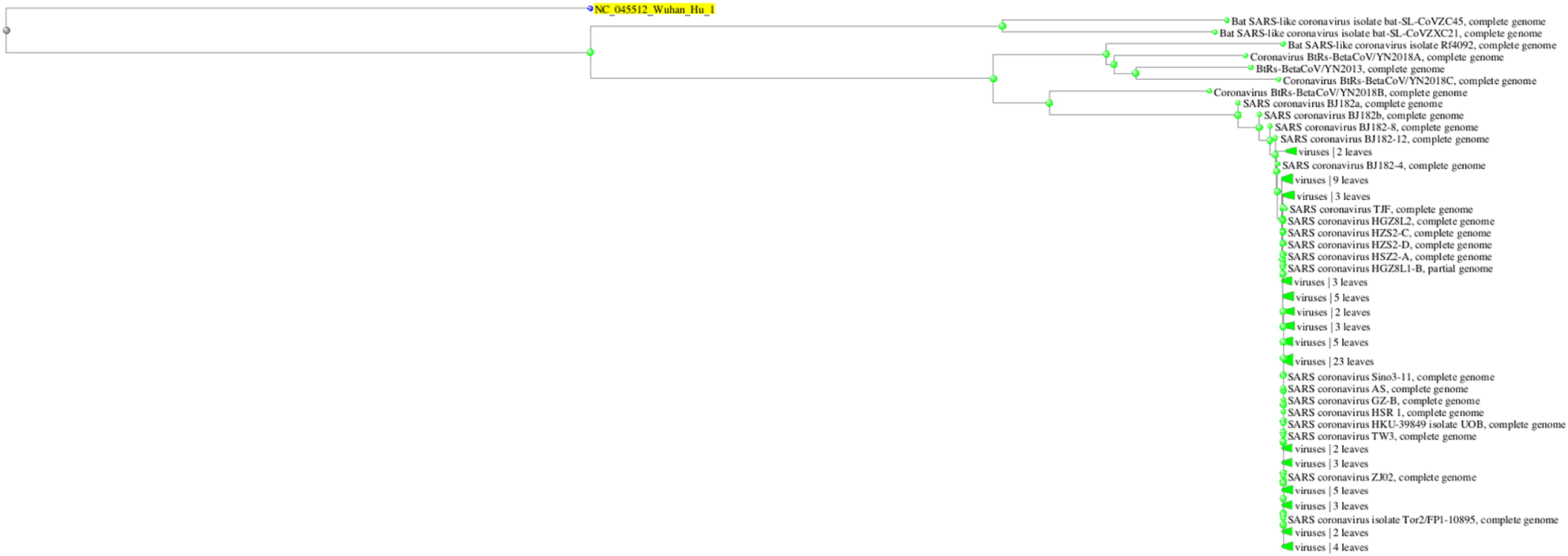
Neighbor joining distance tree of the BLAST search results of the 2019-nCoV (NC_0455_Wuhan_Hu_1) sequence in the genomic region 10901-22830 of the alignment

Our study suggests that the new corona virus (2019-nCoV) is not a mosaic and it is most closely related with the BatCoV RaTG13 detected in bats from Yunnan Province [21]. The levels of genetic similarity between the 2019-nCoV and RaTG13 suggest that the latter does not provide the exact variant that caused the outbreak in humans, but the hypothesis that 2019-nCoV has originated from bats is very likely. On the other hand, there is evidence for discordant phylogenetic relationships between 2019-nCoV and RaTG13 clade with their closest partners the Bat_SARS-like coronavirus sequences. In accordance with previous analysis (http://virological.org/t/ncovs-relationship-to-bat-coronaviruses-recombination-signals-no-snakes/331), bat SARS-like coronavirus sequences cluster in different position in the tree, suggesting that they are recombinants, and thus that the 2019-nCoV and RaTG13 are not [28, 29]. One previous study based on codon usage analyses suggested that the spike protein of 2019-nCoV might have originated from one yet-unknown unsampled coronavirus through recombination [28]. Codon usage analyses can resolve the origin of proteins with deep ancestry with insufficient phylogenetic signal or invented de novo. The recently published bat coronavirus sequence however provides strong phylogenetic information to resolve the origin of the Spike protein as well as the rest of the genome suggesting a uniform ancestry across the genome. We have previously shown that phylogenetic discordance in deep relationships of coronaviruses is common and can be explained either by ancient recombination event or altered evolutionary rates in different lineages, or a combination of both [29]. Our study rejects the hypothesis of emergence as a result of a recent recombination event. Notably, the new coronavirus provides a new lineage for almost half of its genome, with no close genetic relationships to other viruses within the subgenus of sarbecovirus. This genomic part comprises also half of the spike region encoding a multifunctional protein responsible also for virus entry into host cells[30, 31]. The unique genetic features of 2019-nCoV and their potential association with virus characteristics and virulence in humans remain to be elucidated.

